# Generation–Establishment Tradeoffs Shape the Temporal Window of Recombinant Evolution

**DOI:** 10.64898/2026.04.07.717007

**Authors:** S.J. Anthony, H.L. Wells, U. Mitra, P.K. Newton

## Abstract

Recent viral coinfection experiments show that recombinant genomes are generated readily and depend strongly on infection timing and order, yet only a small fraction give rise to persistent lineages. We develop a hybrid deterministic–stochastic framework that resolves this discrepancy by coupling density-dependent recombinant generation with stochastic establishment of rare lineages. The resulting hazard of successful lineage formation is generically non-monotonic, increasing with parental abundance through enhanced generation while decreasing under competitive suppression, and exhibits a unique interior maximum in parental-density space. As parental populations evolve, their trajectory across this hazard landscape defines a sharply localized temporal window of evolutionary opportunity. These results reveal a general principle: evolutionary success is determined not only by intrinsic fitness, but by when variants arise within a dynamically changing ecological context.

## I. INTRODUCTION

Rare stochastic events often determine long-term evolutionary outcomes. In viral populations, recombinant genomes are continually generated during coinfection [1], yet only a small fraction establish persistent lineages, with most going extinct shortly after birth even when intrinsically advantageous [1, 2]. Understanding recombinant success therefore requires coupling deterministic ecological dynamics with stochastic extinction in finite populations.

Classical invasion theory predicts establishment thresholds from intrinsic growth and competitive interactions [3], while branching-process theory quantifies extinction probabilities for rare lineages in fixed environments [4]. However, viral infections are dynamically evolving systems: parental strains expand and compete in a time-varying environment. The fate of a newly generated recombinant thus depends not only on its intrinsic growth rate, but on the ecological state of the parental background at the moment of creation [5]. Prior work has examined establishment in changing environments [6] and clarified limits of diffusion approximations [4, 7]. Recent coinfection experiments further show that recombination tracks with coinfection dynamics but yields re-combinant genomes at low abundance, indicating that recombinants are born into rarity and that their production depends strongly on infection timing and order [1]. These observations highlight the central role of ecological context in determining recombinant success.

To resolve this discrepancy between recombinant production and lineage establishment, we develop a deterministic–stochastic framework that unifies recombinant generation and survival. Parental strains obey Lotka–Volterra dynamics, while rare recombinants follow a birth–death process with an absorbing boundary at extinction. This coupling yields a time-dependent hazard Λ(*t*) describing the instantaneous rate of successful lineage formation. Because generation increases with parental abundance while establishment decreases under competition, the hazard is non-monotonic and admits a unique interior maximum. As parental populations evolve, their trajectory across this landscape produces a localized temporal window of evolutionary opportunity (Fig. 1).

**FIG. 1.**
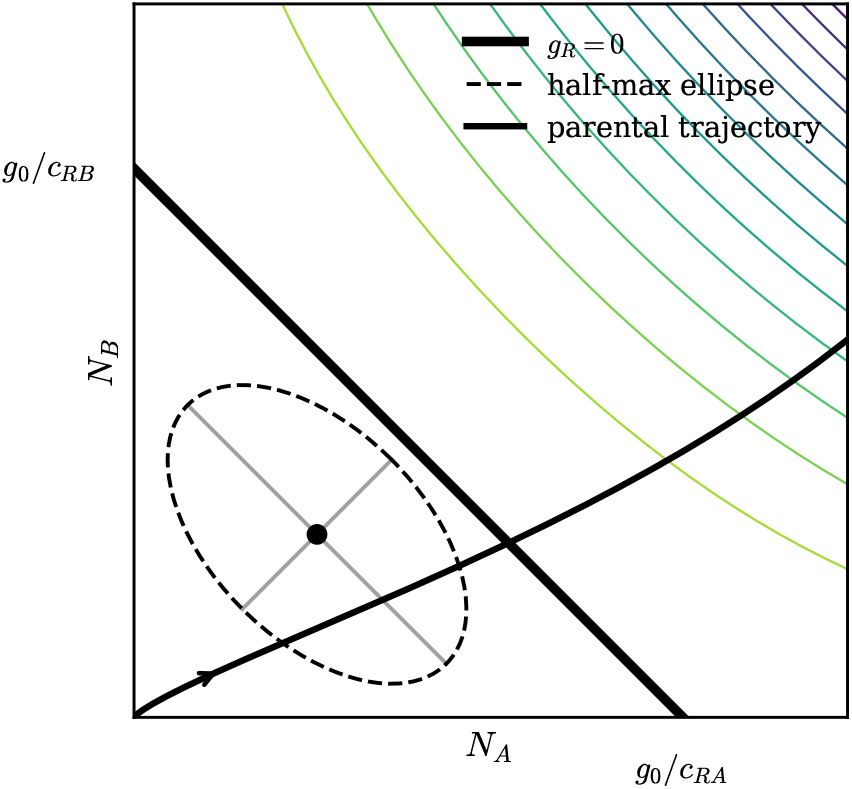
Fate map for recombinant establishment. Contours show the hazard Λ(*N*_*A*_, *N*_*B*_) = *ρN*_*A*_*N*_*B*_ (*g*_0_ − *c*_*RA*_*N*_*A*_ −*c*_*RB*_ *N*_*B*_)*/b*_*R*_. The solid line denotes the invasion boundary *g*_*R*_ = 0 (*g*_*R*_ *>* 0 to the left). The dashed ellipse marks the half-maximum contour, and gray lines indicate principal curvature directions. A parental trajectory highlights the temporal window of maximal establishment.

We derive analytic expressions for extinction probabilities, optimal parental densities, and the timing of establishment, and show that the interior maximum persists under heterogeneous recombinant fitness and saturating coinfection. These results establish a general principle: when generation increases with density while survival decreases under competition, a non-monotonic hazard selects a narrow temporal window in which rare variants can successfully establish.

## II. MODEL: DETERMINISTIC–STOCHASTIC STRUCTURE

We model viral recombination as a coupled deterministic–stochastic system with an explicit separation of scales. Parental strains evolve deterministically due to their large population sizes, whereas newly created recombinant lineages are initially rare and, therefore, governed by stochastic birth–death dynamics. This asymmetry reflects the disparity between macroscopic parental populations and the vanishing abundance of nascent recombinants.

### A. Parental dynamics

Let *N*_*A*_(*t*) and *N*_*B*_(*t*) denote the abundances of parental strains A and B. We adopt a well-mixed competitive Lotka–Volterra description [3],

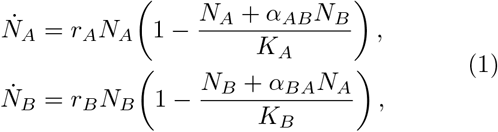

where *r*_*i*_ > 0 are intrinsic growth rates, *K*_*i*_ > 0 carrying capacities, and *α*_*ij*_ > 0 competition coefficients. Because *N*_*A*_ and *N*_*B*_ are assumed large (compared with the recombinant abundances), demographic fluctuations are negligible at this scale.

### B. Recombinant creation

Recombinant genomes are generated during parental coinfection. Creation events are modeled as a non-homogeneous Poisson process [8] with instantaneous rate

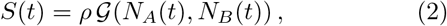

where *ρ* is a recombination efficiency parameter. For low densities, 𝒢 (*N*_*A*_, *N*_*B*_) ∼ *N*_*A*_*N*_*B*_. To incorporate target-cell limitation [9] or mechanisms of competitive suppression (e.g. superinfiction exclusion [1, 10]), a saturating form such as

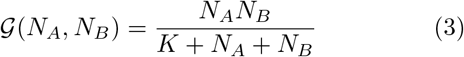

is used.

### C. Recombinant growth and competitive suppression

Each newly created recombinant lineage evolves stochastically. If *N*_*R*_ denotes the abundance of a single lineage, its per-capita birth rate is b_*R*_, while its per-capita loss rate is

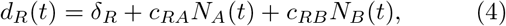

(*c*_*RA*_ > 0 and *c*_*RB*_ > 0 are the competition coefficients between *N*_*R*_ and *N*_*A*_, *N*_*B*_ respectively) so that the net per-capita growth rate is

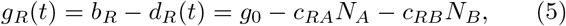

where *g*_0_ ≡*b*_*r*_ −*δ*_*R*_ is the net intrinsic growth when *N*_*A*_ = *N*_*B*_ = 0. Because recombinant lineages are initially rare (*N*_*R*_ ≪ *N*_*A*_, *N*_*B*_), we neglect their feedback on parental dynamics during the establishment phase.

The present analysis concerns the rare regime in which stochastic extinction dominates.

### D. Hazard structure

Recombinant evolution is governed by two distinct processes: (i) the generation of new recombinant lineages at rate *S*(*t*), and (ii) the stochastic establishment of each lineage against demographic extinction. A lineage created at time *t* establishes with probability *P*_est_(*t*)—the eventual survival probability of a single newly created lineage, derived below in Sec. III.

Combining these processes, the rate of successful lineage formation is

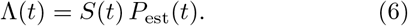

We refer to Λ(*t*) as the hazard [8]: it combines the rate of recombinant generation with the probability of survival, thereby quantifying the rate at which successful recombinant lineages emerge. As a function of parental abundances, Λ(*t*) defines a scalar field over (*N*_*A*_, *N*_*B*_) that governs the timing and likelihood of success.

## III. EARLY-TIME STOCHASTIC ESTABLISHMENT

Newly created recombinant lineages begin at vanishing abundance. We employ a quasi-static (adiabatic) approximation in which parental populations are treated as locally constant during the early stochastic establishment phase (timescale ∼1/*g*_*R*_) of a recombinant lineage. Here, “local” refers to the instantaneous ecological state: establishment probabilities are computed conditional on the parental abundances at the time of recombinant creation, which are treated as approximately constant over this short timescale. Because establishment occurs over a few replication cycles while ecological dynamics evolve more gradually (timescale ∼1/*r*), the hazard structure and its interior maximum are robust to moderate deviations from strict timescale separation.

### A. Branching-process approximation

A lineage founded at time *τ* evolves as a continuous-time birth–death process with per-capita birth rate *b*_*R*_ and death rate (Eq. (4)), with net per-capita growth rate (Eq. (5)).

This implies for a single founder, the eventual extinction probability is the classical branching-process result [8]

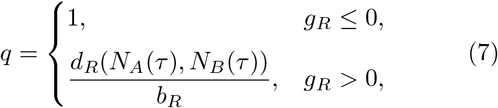

so the establishment probability (1 − *q*) is

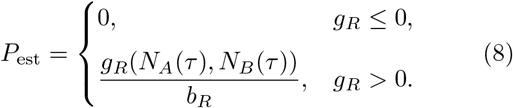

Therefore, positive deterministic growth does not imply a certain success; the probability of establishment scales linearly with the effective growth margin *g*_*R*_(*N*_*A*_(*τ*), *N*_*B*_(*τ*)) at creation (see Supplemental Sec. I A).

### B. Diffusion and Fokker–Planck formulation

The same rare-lineage limit admits a diffusion approximation [4, 7] (see Supplemental Sec. I B)

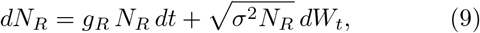

with *σ*^2^ ≈*b*_*R*_ + *d*_*R*_, and absorbing boundary at *N*_*R*_ = 0. *W*_*t*_ is Brownian motion. The corresponding Fokker–Planck (FP) equation for the probability density *p*(*N*_*R*_, *t*) is

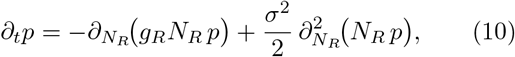

with p(0, t) = 0 enforcing extinction as an absorbing state.

The backward Fokker–Planck equation for the eventual extinction probability [4, 7] yields solution *u*(*N*_*R*_) = exp(−2*g*_*R*_*N*_*R*_/*σ*^2^) for *g*_*R*_ > 0. For a single founder (*N*_*R*_ = 1) and small *g*_*R*_, the exponential expands to *P*_est_ = 1 −*u*(1) ≈2*g*_*R*_/*σ*^2^, which agrees with the classical branching result P_est_ ≈*g*_*R*_/*b*_*R*_ after the usual mapping *σ*^2^ ≈2*b*_*R*_ (since *σ*^2^ ≈*b*_*R*_ + *d*_*R*_ and *d*_*R*_ ≈*b*_*R*_ in the small-drift limit).

## IV. HAZARD LANDSCAPE AND TEMPORAL WINDOW

Combining recombinant creation (Eq.(2)) with stochastic establishment, (Eq.(8)), yields

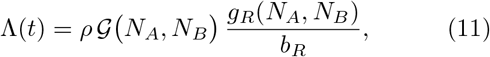

for *g*_*R*_(*N*_*A*_, *N*_*B*_) > 0 (and Λ = 0 otherwise). Because generation 𝒢 increases with parental abundance while *g*_*R*_ decreases under competitive suppression, Λ is non-monotonic.

For symmetric densities *N*_*A*_ = *N*_*B*_ = *N* and bilinear generation 𝒢= *N* ^2^,

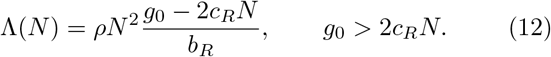

Maximization gives

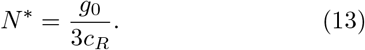

The deterministic invasion boundary *g*_*R*_ = 0 yields *N*_inv_ = *g*_0_/(2*c*_*R*_) in the symmetric case, so *N*^*^ < *N*_inv_. Geometrically, the invasion boundary defines a straight line in parental-density space, whereas the multiplicative generation factor *N*_*A*_*N*_*B*_ produces a curved hazard surface with an interior maximum (Fig. 1).

More generally (see Supplemental Sec. II),

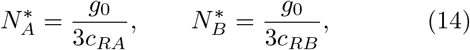

so that 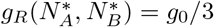. The hazard admits a single interior maximum within the invasion region, so the optial ecological state is uniquely determined. Maximal establishment therefore occurs when increasing generation and decreasing survival are balanced, with the recombinant retaining one-third of its intrinsic growth advantage (Fig. 1).

Viewed over parental-density space (*N*_*A*_, *N*_*B*_), Λ defines a scalar fate map whose level sets partition ecological states by instantaneous establishment rate. Near the maximum, Λ is locally quadratic and its half-maximum contour is an ellipse whose principal axes reflect competitive asymmetry. As the asymmetry increases, the optimum moves towards the axis of weaker suppression and the peak becomes a narrow ridge (see Supplemental Sec. III).

Because parental populations evolve according to Eq. (1), the hazard becomes time-dependent, Λ(*t*) = Λ(*N*_*A*_(t), *N*_*B*_(*t*)), obtained by evaluating the quasi-static hazard along the deterministic parental trajectory. The cumulative probability that at least one recombinant lineage has successfully established by time *T* is

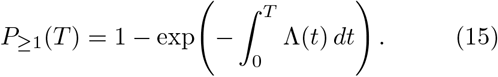

Figure 2 illustrates both the instantaneous hazard Λ(*t*) and the resulting cumulative probability *P*_≥1_(*T*) for unsaturated and saturating recombination models. The marked times correspond to the maxima of the hazard curves, as indicated by the vertical lines in Fig. 2.

**FIG. 2.**
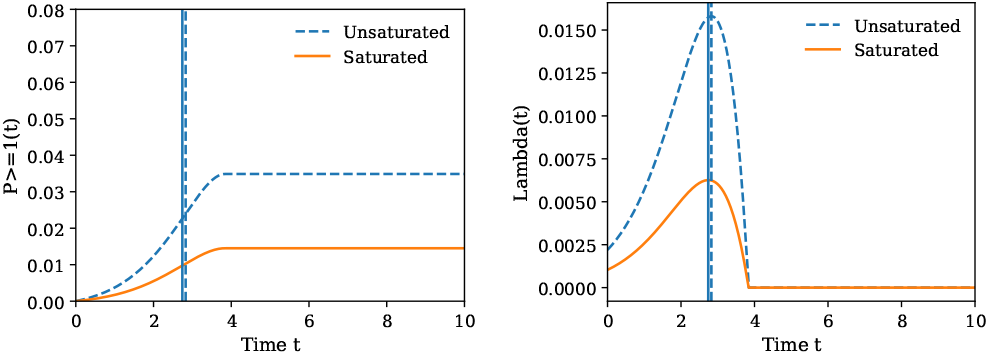
Effect of coinfection saturation on establishment dynamics. (a) Cumulative probability of at least one successful recombinant lineage, *P*_≥1_(*t*) [Eq. (15)], for unsaturated (dashed) and saturating (dash–dotted) recombination. Vertical lines mark the time *t*^*^ of maximal hazard. (b) Corresponding hazard Λ(*t*) along the parental trajectory. Saturation suppresses late-time generation, shifts the peak earlier, and reshapes the temporal window of opportunity.

While *P*_≥1_(*T*) increases monotonically, the optimal time *t*^*^ corresponds to when newly generated recombinants are most likely to successfully establish. This occurs when the hazard 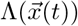 is maximized along the parental trajectory 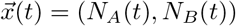, and satisfies

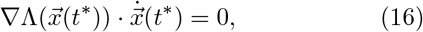

i.e., the trajectory is tangent to a level set of Λ. For symmetric logistic growth, this yields

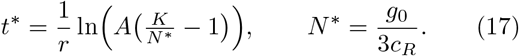

If coinfection saturates (using (3)) the interior maximum persists but shifts toward smaller densities (Fig. 2). With heterogeneous recombinant phenotypes *X*_*R*_ ∼ϕ(*x*) = 𝒩 (*g*_0_, *σ*^2^) (normally distributed r.v. centered at *g*_0_),

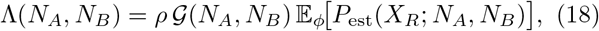

which smooths the peak, but preserves the generation–establishment tradeoff, as shown in Figure 3 for a Gaussian fitness distribution *ϕ*(*x*).

**FIG. 3.**
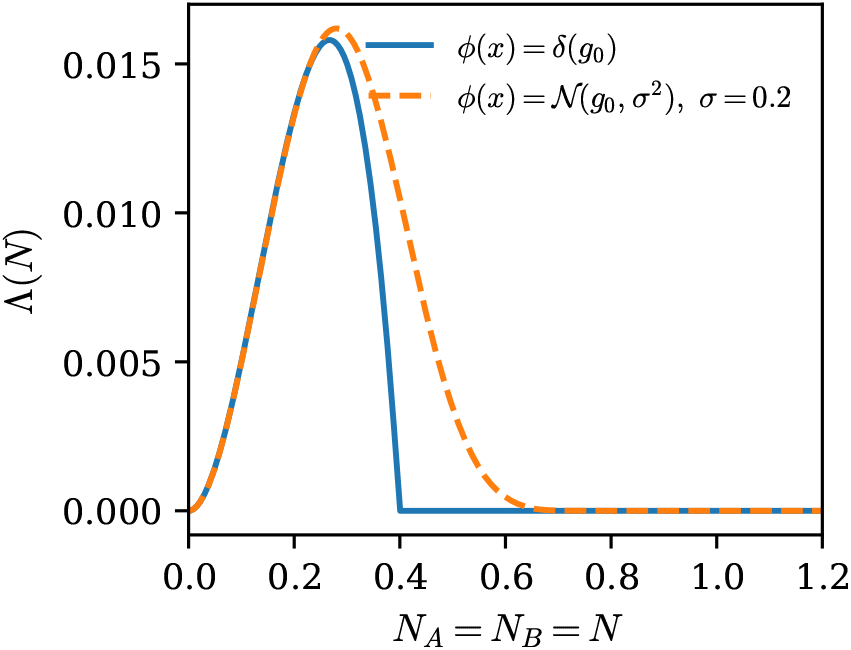
Robustness of the interior maximum to fitness heterogeneity. Hazard Λ(*N*) for symmetric parental densities (*N*_*A*_ = *N*_*B*_ = *N*). Solid: fixed fitness *ϕ*(*x*) = *δ*(*x− g*_0_). Dashed: heterogeneous fitness *ϕ*(*x*) = 𝒩 (*g*_0_, *σ*^2^) (*σ* = 0.2). Averaging over fitness smooths the survival cutoff, broadens the peak, and preserves the interior maximum, demonstrating robustness of the generation–establishment tradeoff.

## V. DISCUSSION

Our results provide a theoretical framework for interpreting recent experiments [1] showing that recombinant genomes are generated readily during coinfection and that their production depends strongly on infection timing and order. These observations indicate that ecological processes govern not only opportunities for recombination but also the subsequent establishment of recombinant lineages, helping to explain why only a subset of recombination events give rise to persistent lineages.

We identify a geometric mechanism linking coinfection dynamics to recombinant establishment: the hazard Λ(*N*_*A*_, *N*_*B*_) combines generation with stochastic survival and is generically non-monotonic, yielding a unique interior maximum within the deterministic invasion region. Invasion criteria alone therefore do not determine evolutionary outcome.

At this optimum, the recombinant retains one-third of its intrinsic growth advantage, reflecting a balance between increasing generation and declining survival under competition. Maximal establishment thus occurs at intermediate ecological states rather than during early expansion or at the invasion boundary. This interior maximum arises under minimal assumptions—monotonic parental growth and density-dependent suppression—ensuring a well-defined optimal ecological state.

As parental populations evolve, they trace trajectories across the hazard landscape, and the temporal window of evolutionary opportunity emerges when the trajectory is tangent to a level set of Λ. Surface curvature governs the robustness of this window, while competitive asymmetry deforms it into ridge-like structures. Saturation and fitness heterogeneity shift the peak but preserve its existence.

More broadly, these results establish a general rare-event principle: when variant generation increases with density while survival decreases under competition, a non-monotonic hazard emerges, and ecological trajectories select a corresponding temporal window of opportunity. This framework yields a directly testable prediction: recombinant yield should depend non-monotonically on infection timing or multiplicity of infection, with a well-defined optimum corresponding to the maximum of Λ along the trajectory (Fig. 2). This can be tested by varying infection timing or multiplicity of infection (MOI) in coinfection experiments and measuring recombinant yield [1].

Evolutionary success is therefore governed not only by intrinsic fitness, but by when variants arise relative to the system’s ecological trajectory.

## SUPPLEMENTAL INFORMATION

### I. BRANCHING AND DIFFUSION PROCESS EXTINCTION FORMULAS

#### A. Extinction probability for the birth–death process

We derive the classical extinction probability for a continuous-time birth–death branching process with percapita birth rate *b*_*R*_ and death rate *d*_*R*_ [8]. Let *q* denote the probability that a lineage starting from a single individual eventually goes extinct.

Over an infinitesimal time interval *dt*, one of three events can occur: (i) no event, with probability 1 −(*b*_*R*_ + *d*_*R*_)*dt*, (ii) death, with probability *d*_*R*_ *dt*, or (iii) birth, with probability *b*_*R*_ *dt*. Conditioning on these events, the extinction probability satisfies

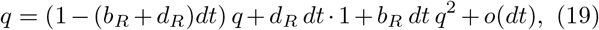

where the factor *q*^2^ arises because a birth event produces two independent lineages, each of which must go extinct. Expanding and subtracting *q* from both sides gives

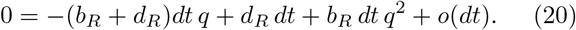

Dividing by *dt* and taking the limit *dt* ⟶ 0 yields

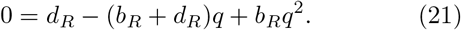

Factoring,

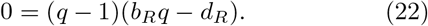

Thus, the extinction probability is

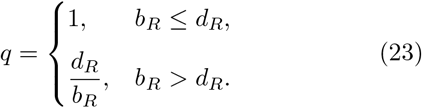

For an initial lineage of size *N* and corresponding extinction probability *q*_*N*_, independence of offspring implies *q*_*N*_ = *q*^*N*^ = (*d*_*R*_/*b*_*R*_)^*N*^, so the establishment probability is

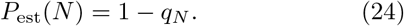

For a single founder (*N* = 1),

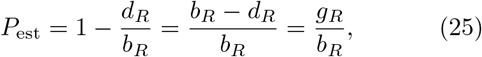

where *g*_*R*_ = *b*_*R*_ − *d*_*R*_ is the net per-capita growth rate.

#### B. Derivation of the diffusion (Fokker–Planck) approximation

To obtain the corresponding diffusion approximation (where *N* becomes a continuous random variable), we start from the discrete birth–death master equation [8] (rate of change = flux in - flux out):

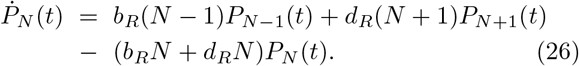

where *P*_*N*_ (*t*) is the probability that a recombinant lineage contains *N* individuals at time *t*.

We then treat *N* as a continuous variable *x* and introduce the probability density *p*(*x, t*) such that *P*_*N*_ (*t*) ≈ *p*(*x, t*) for *x* = *N*. Following the Kramers–Moyal expansion, we write the evolution equation in terms of the jump size Δ*x* = ±1,

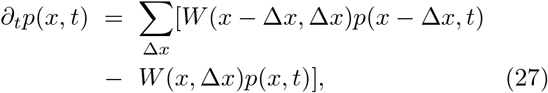

where *W* (*x*, Δ*x*) is the transition rate from *x* to *x* + Δ*x*. For the birth–death process

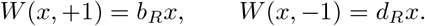

Expanding *p*(*x* ± 1, *t*) in a Taylor series around *x*,

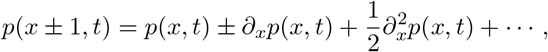

and keeping terms up to second order yields the Kramers–Moyal expansion

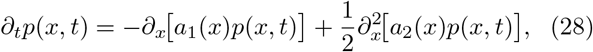

where the first two jump moments are

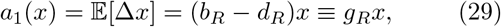

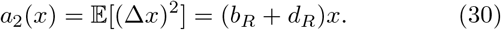

Substituting these moments into Eq. (28) gives the forward Fokker–Planck equation

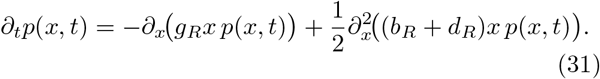

This diffusion equation is equivalent to the Itô stochastic differential equation

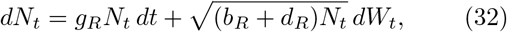

where *W*_*t*_ is standard Brownian motion [7, 8].

#### C. Backward equation and extinction probability

The backward Kolmogorov operator corresponding to Eq. (32) is [8]

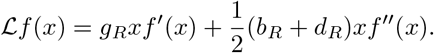

**FIG. S1.**
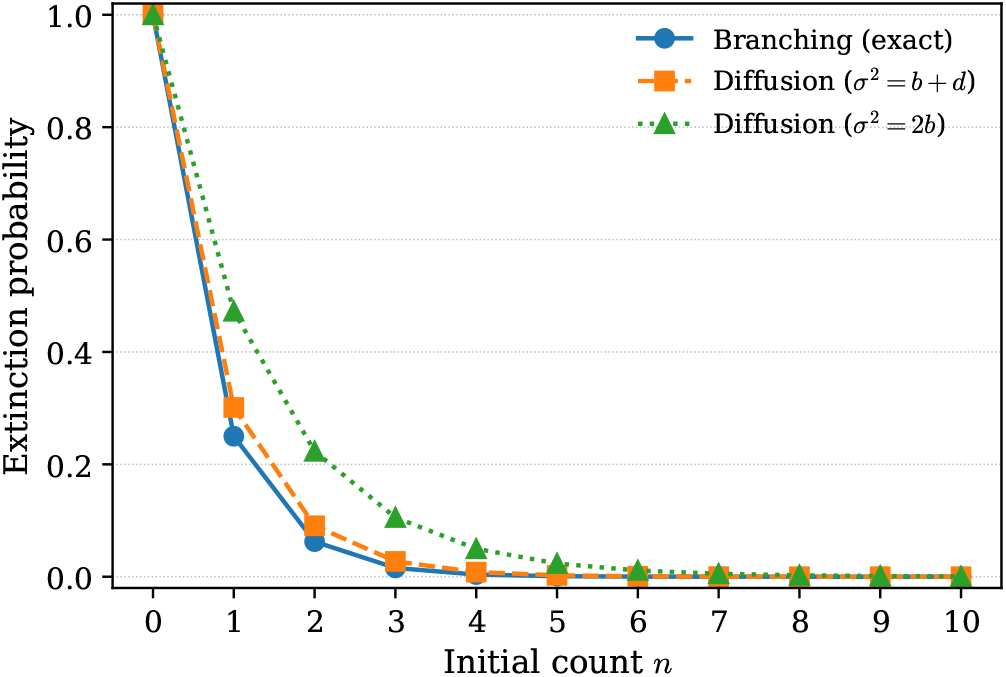
Extinction probability comparison: discrete branching vs. diffusion approximations. Probability of eventual extinction starting from initial count *n* (horizontal axis) computed exactly from the birth–death branching process (*q*^*n*^, solid line, *q* = *d/b*) and from the Fokker–Planck / diffusion approximation *u*(*n*) = exp − 2*gn/σ*^2^ with two noise parametrizations: *σ*^2^ = *b*+*d* (dashed) and *σ*^2^ = 2*b* (dotted). Parameters used: *b* = 2.0, *d* = 0.5 (so *g* = *b* − *d* = 1.5). The diffusion approximations differ noticeably from the exact discrete result for small *n* but converge toward the branching result as *n* increases, illustrating the regime of validity of the continuous approximation and the effect of the heuristic mapping *σ*^2^ *≈* 2*b*.

The eventual extinction probability *u*(*x*) therefore satisfies

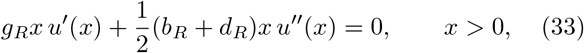

with boundary conditions

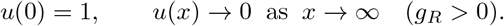

Because each term in Eq. (33) is proportional to *x*, the factor cancels for *x* > 0, leaving a constant-coefficient ODE whose solution is

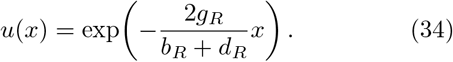

For a single founder (*x* = 1) the establishment probability is

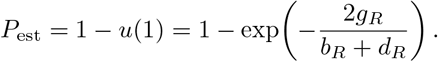

For small *g*_*R*_ this reduces to

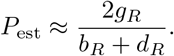

Comparing with the classical branching-process result *P*_est_ ≈ *g*_*R*_/*b*_*R*_ shows that the two approaches agree in the near-critical regime when *b*_*R*_ + *d*_*R*_ ≈ 2*b*_*R*_ (i.e. when *d*_*R*_ ≈ *b*_*R*_) as shown in Fig. S1.

The diffusion equation represents the leading-order truncation of the Kramers–Moyal expansion and is accurate when the probability distribution varies smoothly on the scale of the unit jump. For single-founder extinction probabilities the exact branching-process result is preferable, whereas the diffusion approximation is useful for computing fluxes into the absorbing state, mean extinction times, and corrections for larger initial population sizes [4].

### II. ANALYTIC CALCULATION OF OPTIMAL PARENTAL DENSITIES

Dropping the prefactor *ρ*/*b*_*R*_, define

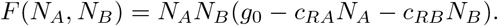

Stationary points satisfy

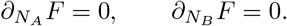

Solving yields the interior optimum

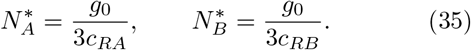

Evaluating the net growth at the optimum gives

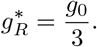

### III. CURVATURE AND HALF-MAX ELLIPSE

Expanding the hazard to quadratic order about the optimum,

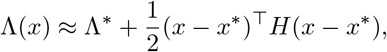

with *A* = −*H* positive definite, the half-maximum contour satisfies

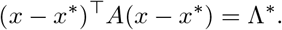

For the explicit hazard,

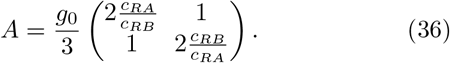

Defining *α* = *c*_*RA*_/*c*_*RB*_, the eigenvalues are

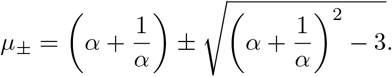

The semi-axis lengths are therefore

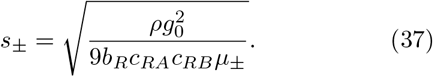

The hazard surface in an asymmetric setting is shown in Fig. S2.

**FIG. S2.**
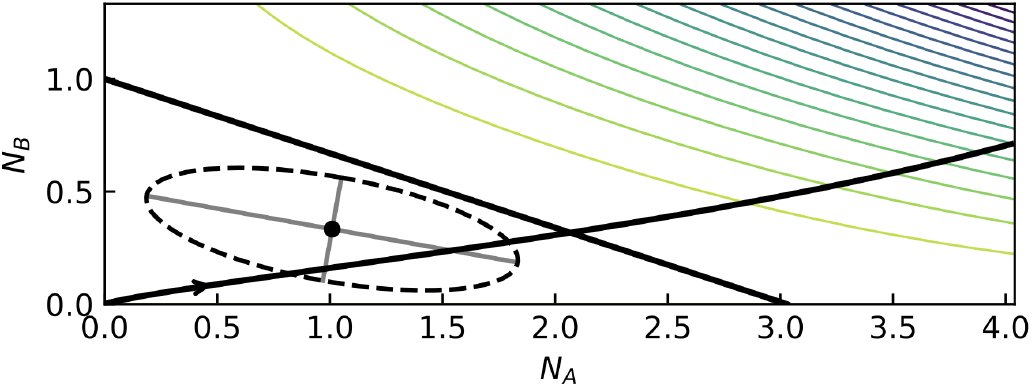
Asymmetric hazard landscape illustrating ridge-like half-maximum contour. Hazard surface Λ(*N*_*A*_, *N*_*B*_) = *ρN*_*A*_*N*_*B*_ (*g*_0_ *− c*_*RA*_*N*_*A*_ *− c*_*RB*_ *N*_*B*_)*/b*_*R*_ for asymmetric competitive suppression with *α* = *c*_*RA*_*/c*_*RB*_ = 0.33. The solid line denotes the deterministic invasion boundary *g*_*R*_ = 0, which intersects the axes at *N*_*A*_ = *g*_0_*/c*_*RA*_ and *N*_*B*_ = *g*_0_*/c*_*RB*_. The dashed curve shows the 50% maximum contour around the interior optimum 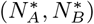. Because suppression by strain *A* is weaker than by strain *B*, the contour becomes strongly elongated along the *N*_*A*_ direction, forming a ridge-like structure in parental-density space.

